# A complete approach for circRNA therapeutics from purification to lyophilized delivery using novel ionizable lipids

**DOI:** 10.1101/2024.10.28.620632

**Authors:** Esther Broset, Verónica Lampaya, Ana Larraga, Víctor Navarro, Alexandre López, Diego de Miguel, Álvaro Peña, Juan Martínez-Oliván, Diego Casabona

**Affiliations:** Certest Pharma, Certest Biotec S.L, Polígono Industrial Río Gallego II, Calle J, 1, San Mateo de Gállego, 50840, Spain

## Abstract

Circular RNA (circRNA) has gained significant attention as a potential therapeutic tool due to its remarkable stability and resistance to degradation by exonucleases. However, scalable and efficient methods for purification and delivery remain critical challenges that must be addressed. In this study, we developed and evaluated an optimized affinity chromatography method using Oligo(dT) columns for the purification of circRNA, achieving high yield and purity comparing with high-performance liquid chromatography. Additionally, we investigated the *in vivo* efficacy of circRNA-Oligo(dT) encapsulated in lipid nanoparticles (LNPs) formulated with emerging STAAR ionizable lipids, including CP-LC-0867 and CP-LC-0729. Our results showed that LNPs formulated with CP-LC-0867 consistently produced higher protein expression compared to SM-102, with sustained luciferase activity observed over a 14-day period. Furthermore, we assessed the lyophilization potential of LNP-circRNA-Oligo(dT) using CP-LC-0729 to extend shelf life and eliminate the need for ultra-low temperature storage. Remarkably, the lyophilized LNPs exhibited no significant differences in protein expression compared to their non-lyophilized counterparts, demonstrating that lyophilization is a viable strategy for extending the storage and transport of circRNA therapies. These findings underscore the potential of optimized new ionizable lipids, improved purification strategies, and lyophilization techniques to enhance the scalability, stability, and practical application of circRNA therapies.

**GRAPHICAL ABSTRACT:** 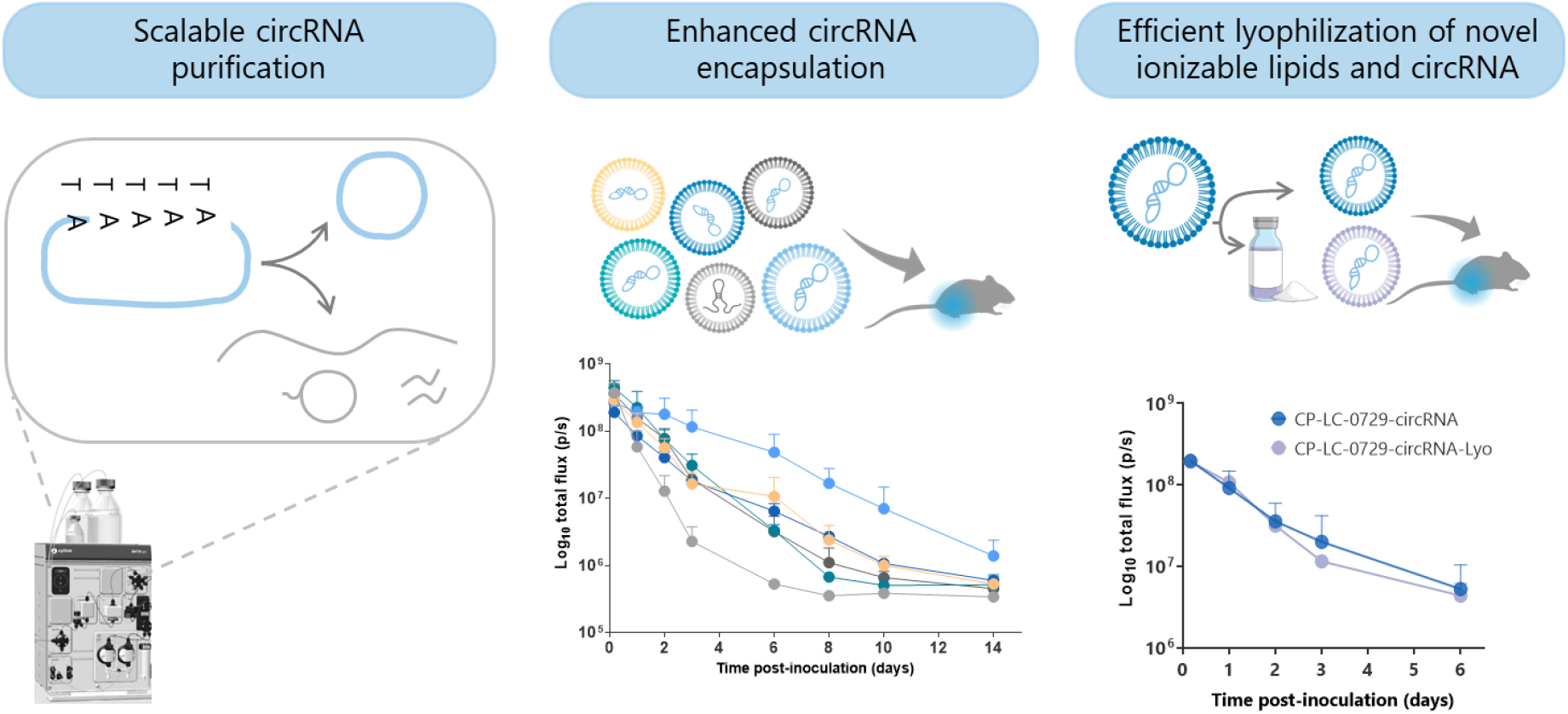

## INTRODUCTION

Circular RNA (circRNA) has emerged as a novel and versatile class of RNA molecules, distinguished by their covalently closed loop structure. This topology grants circRNA remarkable stability, as it lacks the 5’ and 3’ ends that make linear RNA susceptible to exonuclease-mediated degradation. Furthermore, the use of internal ribosome entry site (IRES) in circRNAs allows then to be translated (1) which has increased the attention they have received in various fields, including gene therapy (2), vaccine development (3, 4), and synthetic biology (5). Its ability to sustain prolonged protein expression makes circRNA an attractive alternative to traditional linear RNA (6), particularly in therapeutic applications where long-lasting effects are desired without the risks associated with genomic integration.

The production of circRNA typically involves *in vitro* transcription (IVT) with a subsequent circularization and a complex purification a process to separate circular from linear RNA species (7). The circularization process has recently evolve from the use of low effective T4 ligase mediated reactions (8) to the utilization of highly efficient engineered permuted intron-exon (PIE) self-splicing strategy to promote the circularization of RNA molecules (9, 10). However, the efficient purification of circRNA from precursor, introns and nicked RNA remains a significant challenge. Those RNA contaminants can compromise the immunity and functionality of circRNA (11, 12), which is particularly problematic in applications that require high fidelity and minimal interference, such as in RNA vaccines and gene therapies (13). Traditional methods for RNA purification, such as gel electrophoresis (5) and high-performance liquid chromatography (HPLC)(9), have been used to isolate circRNA. Nevertheless, RNA yields from these methods are typically less than 1% of the input RNA (2, 9), making large-scale circRNA production economically challenging.

Lipid nanoparticle (LNP) system is one of the more effective for *in vivo* delivery of circRNA-based therapies. LNPs are typically composed of four key components: ionizable lipids, phospholipids, cholesterol, and polyethylene glycol (PEG)-lipids. Among these, ionizable lipids are crucial for the efficient release of RNA cargo into the cytoplasm. These ionizable lipids acquire a positive charge in acidic environments, such as within endosomes, facilitating the release of RNA into the cytoplasm. Economically, ionizable lipids represent a significant cost driver in RNA therapies due to their specialized function and the limited number of clinically approved options (14). Currently, MC3, SM-102, and ALC-0315 are among the most widely studied ionizable lipids for clinical use in RNA delivery. MC3, as part of an older generation of lipids, has certain limitations, while SM-102 and ALC-0315 face challenges related to restricted licensing or exorbitant fees, which further complicate access and increase the expense of RNA-based treatments (15). To make circRNA therapies more affordable, the development of new, cost-effective ionizable lipids is essential.

Additionally to the lipid components, the physical and chemical stability of LNP formulations represent another significant challenge for their widespread application (16). Current RNA-LNP therapies, including the approved mRNA vaccines Comirnaty and Spikevax, require cryogenic preservation and transportation between −80 °C to −60 °C for Comirnaty and at −20 °C for Spikevax (17). The stringent storage requirements are primarily due to the intrinsic sensitivity of mRNA to environmental factors such as oxygen, enzymatic degradation, and fluctuations in pH. Additionally, complex interactions between the lipid components and degradation by-products from oxidation and hydrolysis further compromise the stability of mRNA. As water, oxygen, and lipids are included on mRNA-LNP formulations, ensuring high stability in liquid mRNA-LNP systems has historically presented some difficulties (18).

Enhancing the stability of mRNA-LNP therapies has shown promise through lyophilization, a process that removes water by sublimation under vacuum at low temperatures. (19, 20). One study reported successful lyophilization of mannose-modified LNPs containing SM-102 as the ionizable lipid, resulting in a durable immune response against rabies and SARS-CoV-2 viruses (21). While lyophilization has been primarily developed for mRNA-based LNPs, there is limited research on its application to circRNA-LNP formulations. Consequently, developing a circRNA-LNP platform that incorporates novel ionizable lipids capable of withstanding the lyophilization process will be crucial for making these therapies more accessible and practical for public health applications.

In this study, we introduce a complete circRNA platform designed to overcome the economic and industrial challenges associated with circRNA-LNP-based therapies. To address the critical issue of low purification yields, we propose an innovative purification method utilizing poly(A) affinity chromatography. This approach significantly enhances purification efficiency, achieving yields comparable to those of mRNA, thus rendering the process feasible for industrial-scale production. Furthermore, we address the challenge of ionizable lipid selection, a key component in LNP formulations, by exploring novel lipid candidates’ performance with circRNA (22, 23). Finally, we integrate a lyophilization technique tailored to circRNA-LNP formulations (24), ensuring the practicality and accessibility of these therapies for widespread health applications.

## MATERIALS AND METHODS

### Template plasmid construction for circRNA

The DNA template for circRNA synthesis was constructed by cloning the Anabaena 3.0 ribozyme, the internal ribosome entry site IRES-CVB3, both as previously described (9), and a firefly luciferase coding sequence optimized for human expression into a pUC-based plasmid under the control of a T7 promoter. Gene synthesis, cloning, and plasmid preparations were outsourced to Genscript.

Subsequently the plasmid was linearized with the BspQI restriction enzyme (Hongene) at 50°C for 2 hours. The linearized plasmid was then purified using the Wizard® SV gel and PCR clean-up system kit (Promega) following the manufacturer’s instructions.

### Synthesis of RNA and enrichment of circRNA

Modified linear mRNA and unmodified circRNA was synthesized by *in vitro* transcription (IVT) using T7 RNA polymerase (Hongene). The transcription reaction was set up with 1 μg of linearized plasmid DNA, 100 U of T7 RNA polymerase, 5 mM rNTPs, and 1x transcription buffer in a total volume of 20 μL. IVT reaction was incubated at 37°C for 3 hours. Following the IVT reaction, the DNA template was digested with 4 U of DNase I (Hongene) at 37°C for 15 minutes. For circRNA synthesis optimization, we systematically varied the IVT incubation times (1, 2, 3, 5, 7, or 16 hours) and adjusted the concentration of Mg^2+^ in the transcription buffer to 5, 10, 16.5, or 20 mM.

To enrich the circRNA, the IVT product was denatured at 70°C for 5 minutes and then immediately placed on ice for 3 minutes. Subsequently, GTP was added to a final concentration of 2 mM, and the mixture was incubated at 55°C for 15 minutes. The reaction was then quenched by the addition of 5mM EDTA at pH 8 and storage for its subsequent purification.

The control circular RNA (circRNA-Ctrl) used in this study was sourced from GenScript, where it was purified via high-performance liquid chromatography (HPLC).

The control circular RNA (circRNA-Ctrl) used in this study was obtained from GenScript, where it was purified via high-performance liquid chromatography (HPLC) (31). This control circRNA contains the same elements as the experimental circRNA constructs produced in our laboratory, ensuring consistency in evaluating the effects of various conditions on circRNA functionality and stability (37).

### Purification and identification of circRNA

Resulting RNA transcripts were purified by affinity chromatography using POROS Oligo (dT)25 (ThermoFisher). The columns were equilibrated with 5 column volumes of binding buffer (50mM sodium phosphate, 0.5M NaCl, 5mM EDTA, pH = 7.0) before use. The RNA solution was applied to the equilibrated Oligo dT column. The column was washed with 10 column volumes of wash buffer (50mM sodium phosphate, 5mM EDTA, pH = 7.0) and was eluted by applying 2 column volumes of double-deionized water. The eluate was collected and immediately placed on ice.

Subsequently, a polishing treatment of the RNAs was performed to enrich the circular isoform relative to the other species. Resulting RNA fractions were visualized on a 4% polyacrylamide gel (PAGE) containing 8 M urea and analyzed using the FIJI software.

The circRNA-Kit was purified using the Monarch® Spin RNA Cleanup Kit (50 μg), following the manufacturer’s instructions for optimal RNA recovery.

The quality and size of the resulting RNA samples were evaluated using a Bioanalyzer (Agilent 2100) with the Agilent RNA 6000 Nano kit. The resulting data were processed and evaluated with 2100 Expert Software. RNA quality was further evaluated using high-performance liquid chromatography (HPLC). Samples were processed through a CIMac™ SDVB 0.1 mL analytical reverse-phase column (2 μm channels, Sartorius) connected to an HPLC WATERS e2695. CIMac SDVB column was equilibrated at 65ºC in mobile phase A (50mM TEAA and 7.5% ACN, pH 7.0). Approximately 0.4 μg RNA was injected on the column. After an initial equilibration step (1 min at 100% MPA), a linear gradient was performed, 0–100% B (50mM TEAA in 18% CAN pH = 7.0) in 27 min at a flow rate 1mL min^−1^. The column was eluted with 100% MPC (50mM TEAA in 75% CAN pH = 8.2) for 1 min before re-equilibration in 100% MPA.

### Determination of %dsRNA content

The circRNAs were spotted onto positively charged nylon membranes (Nytran SC, Sigma Aldrich). The membranes were then blocked with 5% (w/v) non-fat dried milk in TBS-T buffer (20mM Tris–HCl, 150mM NaCl, 0.1% [v/v] Tween-20, pH 7.4), and incubated with dsRNA-specific mAb J2 (Jena Bioscience) at 4ºC overnight. Membranes were washed with TBS-T buffer and incubated with HRP-conjugated goat anti-mouse IgG (Abcam). Following additional wash with TBS-T, the membranes were treated with ECL Plus Western blot detection reagent (Amersham). Images were captured using an iBright 750 digital imaging system.

### Synthesis and Characterization of Ionizable Lipids

The ionizable lipids CP-LC-0743, CP-LC-0729, and CP-LC-1254 were synthesized following established protocols based on the Sequential Thiolactone Amine Acrylate Reaction (STAAR) (21). In brief, thiolactone derivatives (0.15 mmol, 1 equiv.) and acrylate (0.15 mmol, 1 equiv.) were dissolved in 300 μL of tetrahydrofuran (THF) at room temperature (RT), followed by the addition of the amine (0.15 mmol, 1 equiv.). After two hours, the THF was removed under vacuum. Notably, CP-LC-0743 was previously referred to as A4B2C1, and CP-LC-1254 was previously designated as CP-LC-0729-04 (21).

For the synthesis of CP-LC-0867, B5 compound (22)(0.15 mmol, 1 equiv.) and 2-ethylhexyl acrylate (0.15 mmol, 1 equiv.) were dissolved in 300 μL of THF at RT, followed by the addition of N,N-dimethylethylenediamine (0.15 mmol, 1 equiv.). After two hours, the THF was removed under vacuum, and the product was purified using a CombiFlash NextGen 300+ with gradient elution, starting from 100% dichloromethane to 50% of an 80/20/1 mixture of DCM/MeOH/NH_4_OH (aq).

All synthesized lipids were analyzed by high-performance liquid chromatography (HPLC) equipped with a charged aerosol detector (CAD) and characterized by mass spectrometry (ThermoFisher ISQ). The resulting theoretical and experimental masses were as follows: CP-LC-0743: theoretical [M+H]+ = 766.25, experimental [M+H]+ = 766.81, CP-LC-0729: theoretical [M+H]+ = 740.63, experimental [M+H]+ = 740.85, CP-LC-1254: theoretical [M+H]+ = 755.63, experimental [M+H]+ = 755.83, CP-LC-0867: theoretical [M+H]+ = 684.57, experimental [M+H]+ = 684.75.

### Lipid nanoparticle (LNP) formulation and characterization

The synthesis of LPNs was performed using a microfluidic method. An ethanolic lipid solution containing the synthetized STAAR ionizable lipids or the commercial SM-102 (BocSCI), DOPE (Corden Pharma), cholesterol (Merk), and DMG-PEG2000 (Cayman) at a molar ratio of 50:10:38.5:1.5 was prepared. This lipid mixture was then combined with an aqueous phase containing mRNA or circRNA diluted in 10 mM citrate buffer (pH 4), achieving an ionizable lipid/RNA weight ratio of 10:1. LNPs were formulated using an INano™ microfluidic device set to a total flow rate of 12 mL/min with an aqueous-to-ethanol volume ratio of 3:1. After LNP formation, ethanol was removed by dialysis (Pur-A-Lyzer™ Midi Dialysis Kit, Sigma).

For LNP characterization the average size, polydispersity (PDI), and zeta potential of LNPs were determined using a Malvern Zetasizer Advance Lab Blue Label (Malvern Instruments Ltd). Encapsulated RNA was then quantified using the Quant-IT® Ribogreen assay (Invitrogen) following the manufacturer’s instructions, and LNP encapsulation efficiency were verified by agarose gel electrophoresis.

The final LNP formulation was adjusted to an mRNA concentration of 100 μg/mL and was subsequently lyophilized and/or stored at 4ºC for subsequent *in vivo* and *in vitro* analysis.

### LNP Lyophilization

Lyophilization was performed following previously reported protocols (24), using a Virtis Genesis Pilot Freeze Dryer. In brief, LNPs were lyophilized with 20% maltose as a lyoprotectant in TRIS buffer. The process included three stages: (1) freezing at −50 °C, (2) primary drying at −10 °C under 180 mTorr pressure, and (3) secondary drying at 40 °C, gradually reducing the pressure to 120 mTorr and finally to 50 mTorr. Upon completion, the vials were backfilled with pure nitrogen to maintain an inert atmosphere, sealed, and stored at various temperatures for stability assessments. For reconstitution, 300 μL of RNase-free water was added to each vial, and the contents were gently swirled until a homogenous suspension was obtained.

### Cell culture and circRNA transfection

The HeLa (DSMZ GmbH) HEK293T (ATCC) and A549 (NIBSC) cell lines were maintained in DMEM with high glucose (Merk), supplemented with 10% Fetal Bovine Serum (Sigma), 1% Penicillin-Streptomycin Solution (GibcoTM) and 2 mM Glutamax (ThermoFisher). All cell lines were grown in 175 cm^2^ flasks.

The day before transfection, cells were detached from culture flasks using trypsinization and subsequently seeded into 96-well plates at a density of 1 × 10^4^ cells per well. Immediately before the transfections, the culture media in each well was replaced with 90 μL of fresh media. For RNA transfection a mixture containing RNA and Lipofectamine MessengerMAX™ (Invitrogen) was pre-incubated in OptiMEM media according to the manufacturer’s instructions. This mRNA-lipofectamine complex was then added to the designated wells in triplicate, achieving a final RNA concentration of 100 ng/well.

For RNA-LNP transfection a suspension of RNA-containing LNPs was added to the designated wells in triplicate to achieve a final RNA concentration of 200,100 and 50 ng/well. The cells were then incubated 24h at 37°C in a 5% CO_2_ atmosphere before luciferase activity quantification.

### Luciferase activity quantification *in vitro*

At 24- or 48-hours post-transfection, the cells were lysed by adding 100 μL of PBS containing 0.1% Triton X-100 to each well and incubating for 10 minutes at room temperature. Following lysis, 98 μL of the cell lysate was transferred to an opaque 96-well white plate. To each well, 100 μL of a buffered d-Luciferin solution (GoldBio) containing 100 mM Tris-HCl (pH 7.8), 5 mM MgCl_2_, 250 μM CoA, and 150 μM ATP was added, resulting in a final d-Luciferin concentration of 150 μg/mL. Luminescence was measured after a 5-minute incubation at room temperature using a FLUOstar Omega plate reader (BMG LABTECH). Cells that had not been treated with any mRNA were used as a negative control to assess background luminescence.

### Administration of LNP-circRNAs in mice

Female BALB/c mice (Janvier), aged 8-10 weeks and weighing 18-23 g, were acclimatized to the experimental facilities for 3-7 days upon arrival. The mice were housed under controlled conditions, maintaining a room temperature of 20-24°C, humidity levels between 50-70%, and a light intensity of 60 lux with a 12-hour light-dark cycle.

For *in vivo* measurement of Firefly Luciferase activity, LNPs containing 1 μg of the specified RNA in a final volume of 30 μL. were administered via intramuscular injection. At the designated times post-injection, mice were anesthetized with 4% isoflurane administered via a vaporizer for induction, and anesthesia was maintained at 1.5% isoflurane. Following anesthesia, D-luciferin (Quimigen) was administered intraperitoneally at a dose of 150 mg/kg. Ten minutes after D-luciferin administration, luminescence images were captured using the IVIS Lumina XRMS Imaging System.

All animal procedures were conducted under Project License PI07/23, approved by the Ethics Committee for Animal Experiments at the University of Zaragoza. The care and use of animals were carried out in accordance with the Spanish Policy for Animal Protection (RD 53/2013), which aligns with the European Union Directive 2010/63/EU on the protection of animals used for scientific and experimental purposes

## RESULTS

### Optimization of *in vitro* transcription conditions for enhanced circRNA production

To optimize our circRNA production platform, we utilized the PIE self-splicing circularization system, which is widely recognized for its efficiency (9). The initial focus was on enhancing *in vitro* transcription (IVT) conditions, particularly incubation time and Mg^2+^ concentration as both are critical determinants of yield and RNA quality (25). For circRNA production, the Mg^2+^ concentration is particularly important, as it is essential not only for the IVT process but also for the PIE-mediated circularization.

Our optimization began by adjusting the incubation time while maintaining a Mg^2+^ concentration of 16.5 mM and an incubation temperature of 37°C. We observed that a 3-hour incubation period resulted in the highest percentage of circRNA isoforms, indicating it as the optimal duration (Figure 1A). We next varied the Mg^2+^ concentration to evaluate its effect on circRNA yield. In line with our previous optimization studies for mRNA IVT (26), we identified 16.5 mM of Mg^2+^ as the optimal concentration for circRNA production (Figure 1B and C). Under these conditions, approximately 60% of the RNA was in the circular isoform, compared to 40% observed with 20 mM and 10 mM of Mg^2+^. To quantify the proportion of circular isoforms, we performed PAGE electrophoresis coupled with image analysis (Figure 1B) and confirmed the results using capillary electrophoresis, which yielded consistent findings (Supplementary Figure 1). Notably, reducing the Mg^2+^ concentration from 10 mM to 5 mM resulted in a complete absence of RNA synthesis. This underscores the critical role of Mg^2+^ concentration in the formation of both mRNA and circRNA.

**Figure 1.**
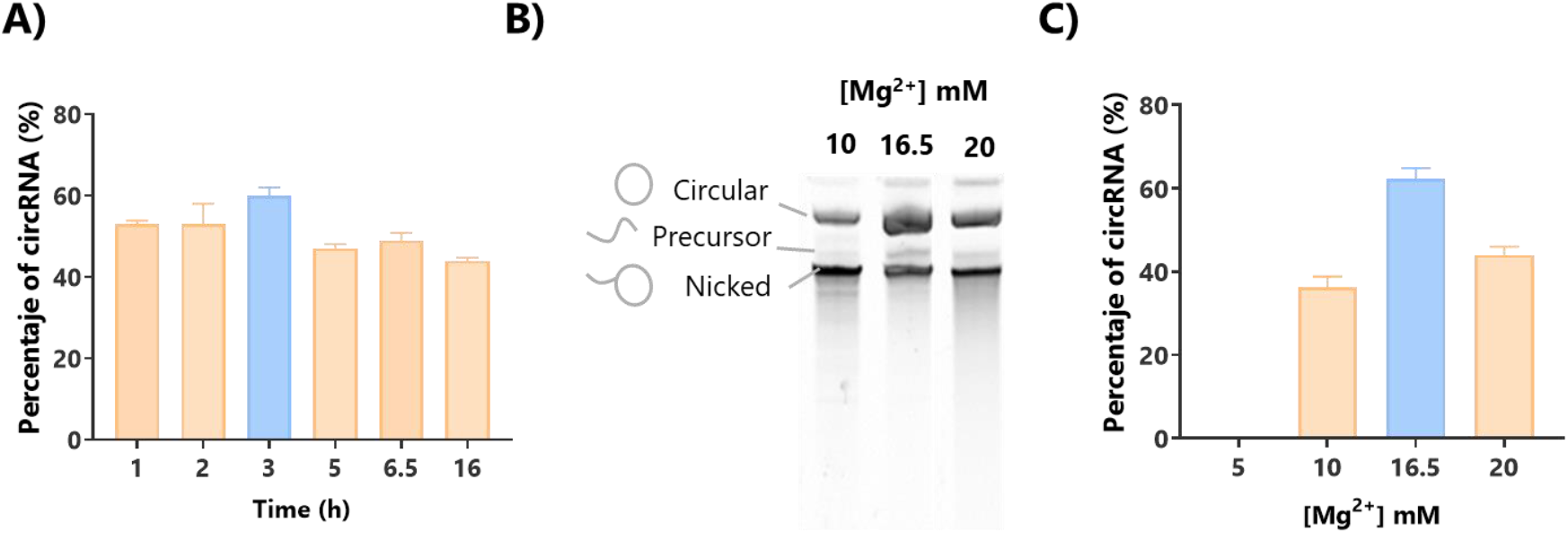
Optimization of incubation time and Mg^2+^ concentration for circRNA production. **(A)** Quantification of circular isoform yield at various incubation times during the *in vitro* transcription (IVT) reaction. **(B)** Polyacrylamide gel electrophoresis (PAGE) analysis of circRNA yield at different Mg^2+^ concentrations (10, 16.5, and 20 mM) used on the IVT reaction buffer. **(C)** Percentage of circular RNA isoform based on PAGE results from panel B quantified by image analysis.

### Scalable circRNA purification using affinity chromatography

After optimizing the IVT conditions, we shifted our focus to the purification steps. Traditionally, circRNA production involves an IVT step followed by circularization with the addition of GTP and Mg^2+^, and a subsequent purification step. The resulting circRNA is then treated with RNase R to eliminate the linear RNA, and then it is further purified. To maximize yield, we chose to bypass the RNase R treatment and the subsequent purification step, directly purifying the circRNA after the circularization step. We leveraged the polyA tail in the circRNA design (9, 27) and utilizing affinity chromatography with Oligo (dT)25 columns (Oligo(dT)) for the purifications. This method was compared to a gold-standard approach commonly used which utilizes silica-based purification. The Oligo(dT) method yielded an average recovery of 18.9 μg of total RNA per μg of IVT input pDNA (Figure 2A). Although this recovery rate is lower than the 55.75 μg per μg of pDNA achieved with the silica-based method (circRNA-kit), it is important to note the reduced presence of non-circular RNA isoforms with the Oligo(dT) method (Figure 2B and C).

**Figure 2.**
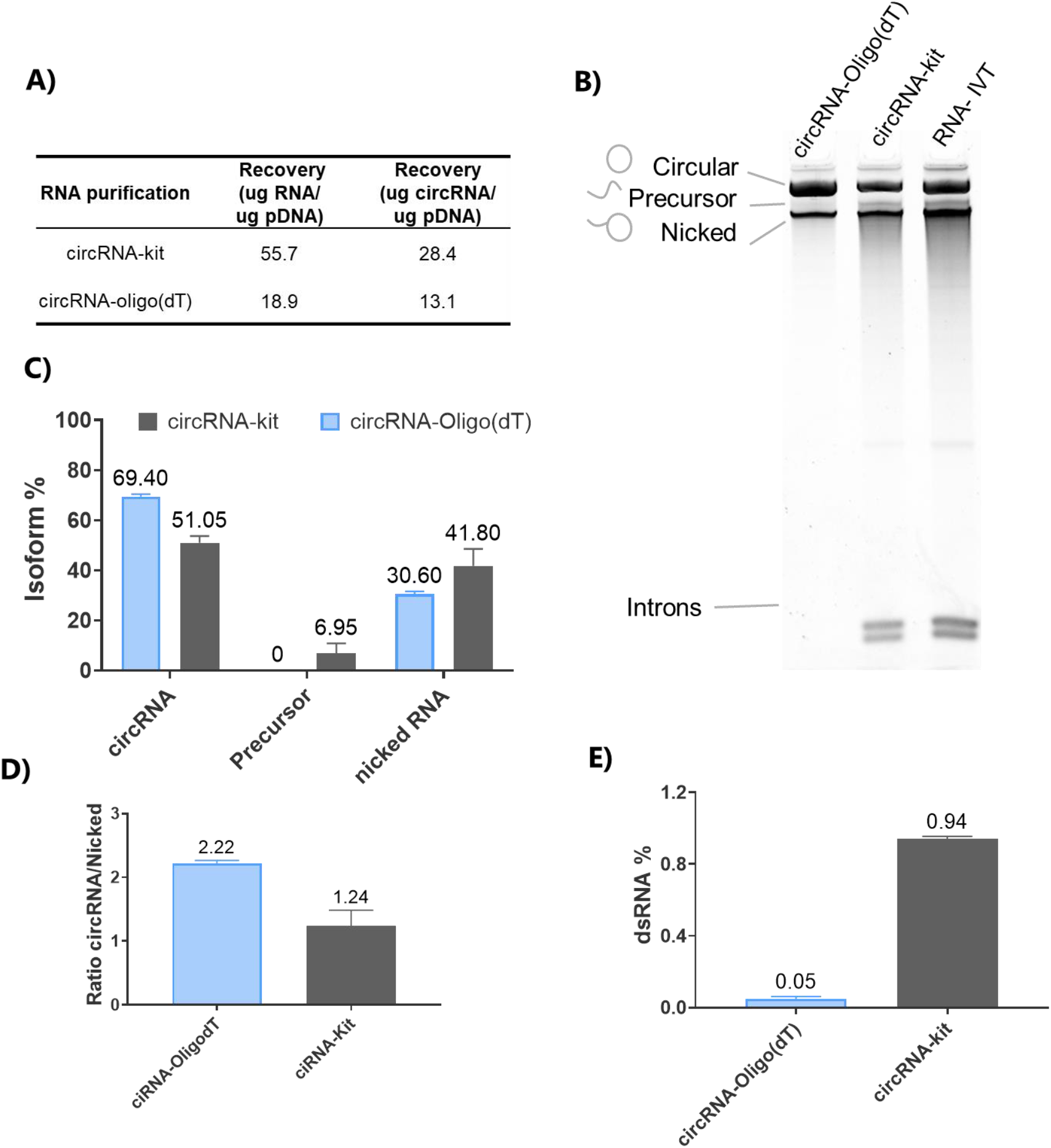
Comparison of circRNA Purification Methods and Quality Assessment. **(A)** Total RNA and circRNA recovery (μg) per μg of IVT input pDNA for Oligo(dT) affinity chromatography compared to the silica-based circRNA-kit method. **(B)** Polyacrylamide gel electrophoresis (PAGE) analysis of RNA before purification (RNA-IVT) and post-purification using either the silica or Oligo(dT) methods. Circular RNA, precursor, nicked, and intronic RNA species are indicated **(C)** Quantification of the percentage of circular RNA isoform based on PAGE results in panel B, determined through image analysis. **(D)** circRNA/nicked RNA ratio from panel C. **(E)** Quantification of dsRNA content through dot-blot analysis, with image-based quantification.

Remarkably, after the Oligo(dT) purification, we observed the complete removal of the intron isoform and linear precursors (Figure 2B and C). This efficient precursor removal justifies omitting the RNase R treatment, thereby eliminating the need for an additional, costly enzymatic step. In contrast, when using the silica-based purification method, intron isoforms and linear precursors remained at levels comparable to those observed immediately after the IVT reaction. Furthermore, when evaluating purity, we found that the circRNA/nicked RNA ratio serves as a reliable purity indicator. Oligo(dT) purification achieved a two-fold higher circRNA/nicked RNA ratio compared to the silica method (Figure 2D).

The presence of dsRNA in RNA therapies has been shown to be detrimental to protein production (28, 29), as it activates the innate cellular response, leading to the elimination of foreign RNA (30). HPLC purification has been demonstrated to eliminate immune activation in IVT mRNA samples (28) and circRNA (9, 13); however, there is no data regarding its effect when using Oligo(dT) purification for circRNA. Therefore, to further assess the quality of the purified circRNA, we quantified the percentage of dsRNA% in the samples (Figure 2E). Our analysis demonstrated that affinity chromatography reduced dsRNA levels to 0.05%, significantly below the 0.5% threshold generally considered acceptable for RNA-based therapies. Notably, compared to circRNA-kit purified using a silica-based method, our circRNA-Oligo(dT) purification resulted in an 18.8-fold reduction in dsRNA content. This substantial decrease emphasizes the capability of Oligo(dT) purification in minimizing dsRNA impurities, which is crucial for enhancing the safety and efficacy of circRNA-based therapies.

### Oligo(dT) purified circRNA exhibits enhanced *in vitro* and *in vivo* translation

We next assessed the translational capacity of circRNA purified by Oligo(dT) affinity chromatography *in vitro*, comparing it to circRNA purified by HPLC (circRNA-Ctrl) as a benchmark (31). Luminescence assays conducted 24 hours post-transfection (Figure 3A) showed that circRNA-Oligo(dT) produced 14-fold higher protein levels compared to the commercial circRNA-Ctrl, and 9-fold compared to a circRNA purified by commercial silica-based capture columns (circRNA-kit) in HeLa cells. Additionally, circRNA-Oligo(dT) exhibited a 2-fold increase in protein production compared to conventional *N*1-methylpseudouridine-modified mRNA (Figure 3A). Similar trends were observed in HEK293T (Supplementary Figure 2) and A549 cells (Figure 3C). Notably, the difference was most pronounced in A549 cells, where circRNA-Oligo(dT) generated 25-fold higher luminescence than circRNA-Ctrl purified using the HPLC method (31) and 9-fold compared to circRNA-kit. This is particularly significant given that A549 cells possess functional Toll-like receptors (TLRs), making them more sensitive to circRNA impurities than HEK293T cells (13). Furthermore, these transfection results align with our dsRNA impurity data, which showed higher levels of impurities in circRNA-kit than in circRNA-Oligo(dT) (Figure 2C).

**Figure 3.**
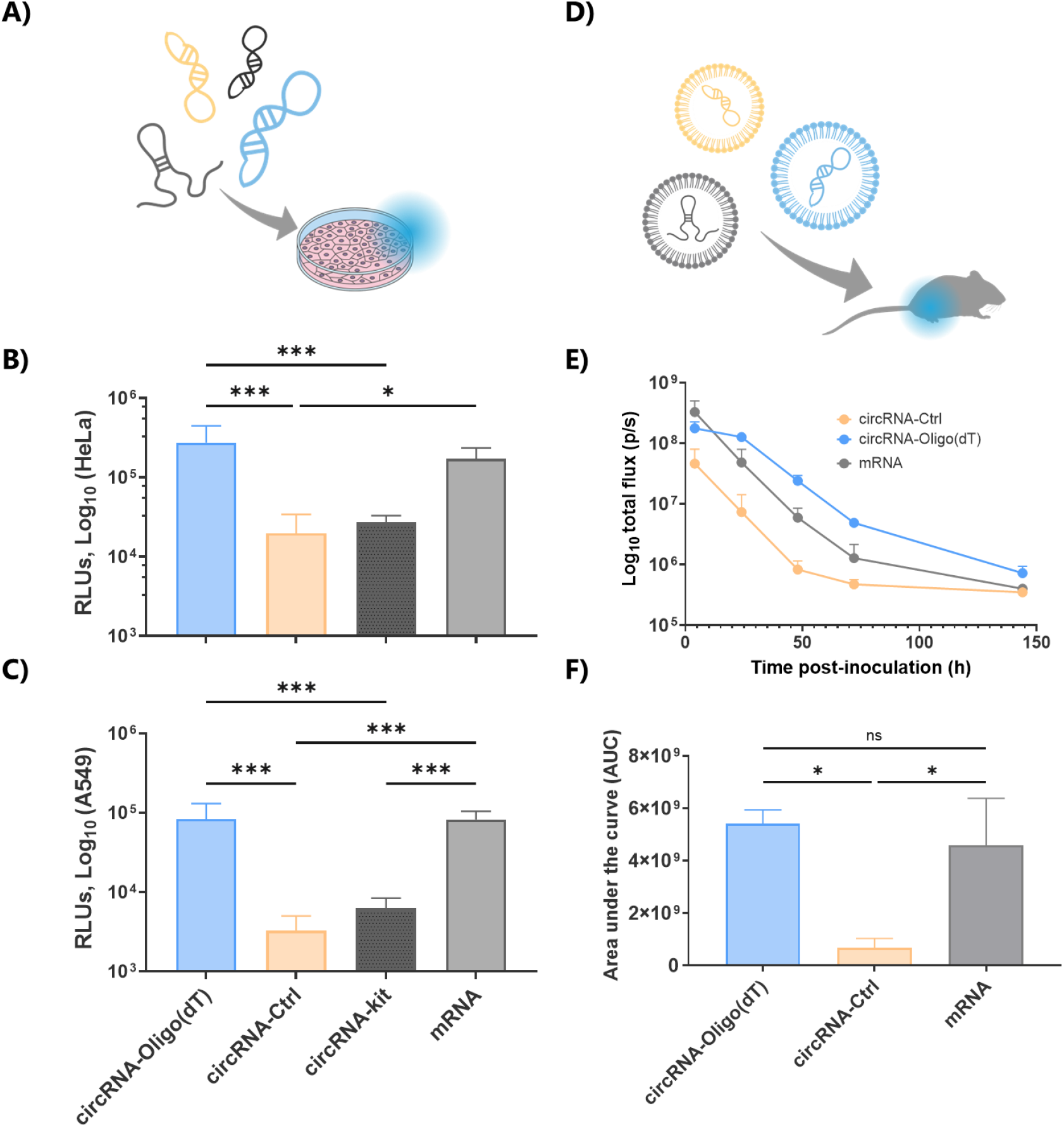
Translation efficacy of circRNA purified by oligo(dT) affinity chromatography. **(A)** Schematic of *in vitro* transfection of RNAs. Oligo(dT)-purified circRNA (circRNA-Oligo(dT)) is represented in blue, commercial control circRNA (circRNA-Ctrl) in yellow, circRNA purified by commercial silica-based capture columns (circRNA-kit) is represented in black and 5-methoxyuridine-modified mRNA (mRNA) in grey. **(B)** HeLa or **(C)** A549 cell lines were transfected with 100 ng/well of circRNA-Oligo(dT), circRNA-Ctrl, circRNA-kit or mRNA. Luminescence was measured 24 hours post-transfection. **(D)** Scheme of the *in vivo* procedure to assess translation efficacy of the different RNAs encapsulated into LNPs. **(E)** Luminescence levels monitored over 144 hours following intramuscular injection of LNPs encapsulating circRNA-Oligo(dT), circRNA-Ctrl, or mRNA in a mouse model. **(F)** Area under the curve (AUC) analysis of cumulative luciferase expression over the 6-day period represented in E. Results are represented as mean ± SD. Statistical significance was determined using one-way ANOVA with Tukey’s post-hoc test (*: *P*-value <0.05**; *P*-value <0.01; ***: *P*-value <0.001; *ns*: *P*-value >0.05).

Given that circRNA is intended for use in *in vivo* therapies, we next assessed the translational capacity of circRNA purified using our new method in a mouse model. We encapsulated circRNA-Oligo(dT), circRNA-Ctrl, and mRNA into conventional LNPs using ionizable lipid SM-102. Following intramuscular administration of RNA-LNPs, luminescence was monitored over 6 days. circRNA-Oligo(dT) showed higher luminescence values than circRNA-Ctrl from time point at 4 h (Figure 3D and E) and also surpassed those of modified mRNA. Area under the curve (AUC) analysis over the 6-day period indicated that circRNA-Oligo(dT) produced a significantly higher cumulative amount of luciferase protein than circRNA-Ctrl (Figure 3F). Although circRNA-Oligo(dT) produced a similar cumulative amount of protein as mRNA, the persistence of the signal was markedly different. By day 3 (72 hours), luminescence from circRNA-Oligo(dT) was 3.7 times higher than that of mRNA, indicating that circRNA-Oligo(dT) may be better suited for therapies requiring sustained protein expression over time.

### circRNA-oligo(dT) encapsulated with novel ionizable lipids shows long lasting protein production in *vivo* and lyophilizability

To further develop an industrially scalable circRNA therapy, it is essential to expand the range of ionizable lipids available for LNP encapsulation. We have developed a broad family of ionizable lipids (22), originally evaluated for mRNA encapsulation and delivery, but with applicability for encapsulating any potential type of nucleic acid. In this study, we focused on evaluating the top three candidates from this family—CP-LC-0729, CP-LC-0743, and CP-LC-1254—specifically for their efficiency in delivering circRNA-Oligo(dT) *in vivo*. In addition, we introduced a novel, unpublished lipid, CP-LC-0867, synthesized using the same STAAR platform, to further expand our evaluation. Each LNP formulation was physically characterized (Supplementary Figure 4A) tested *in vitro* (Supplementary Figure 4B) and evaluated in a mouse model via intramuscular injection to assess its effectiveness in circRNA delivery and subsequent protein expression (Figure 4A).

**Figure 4.**
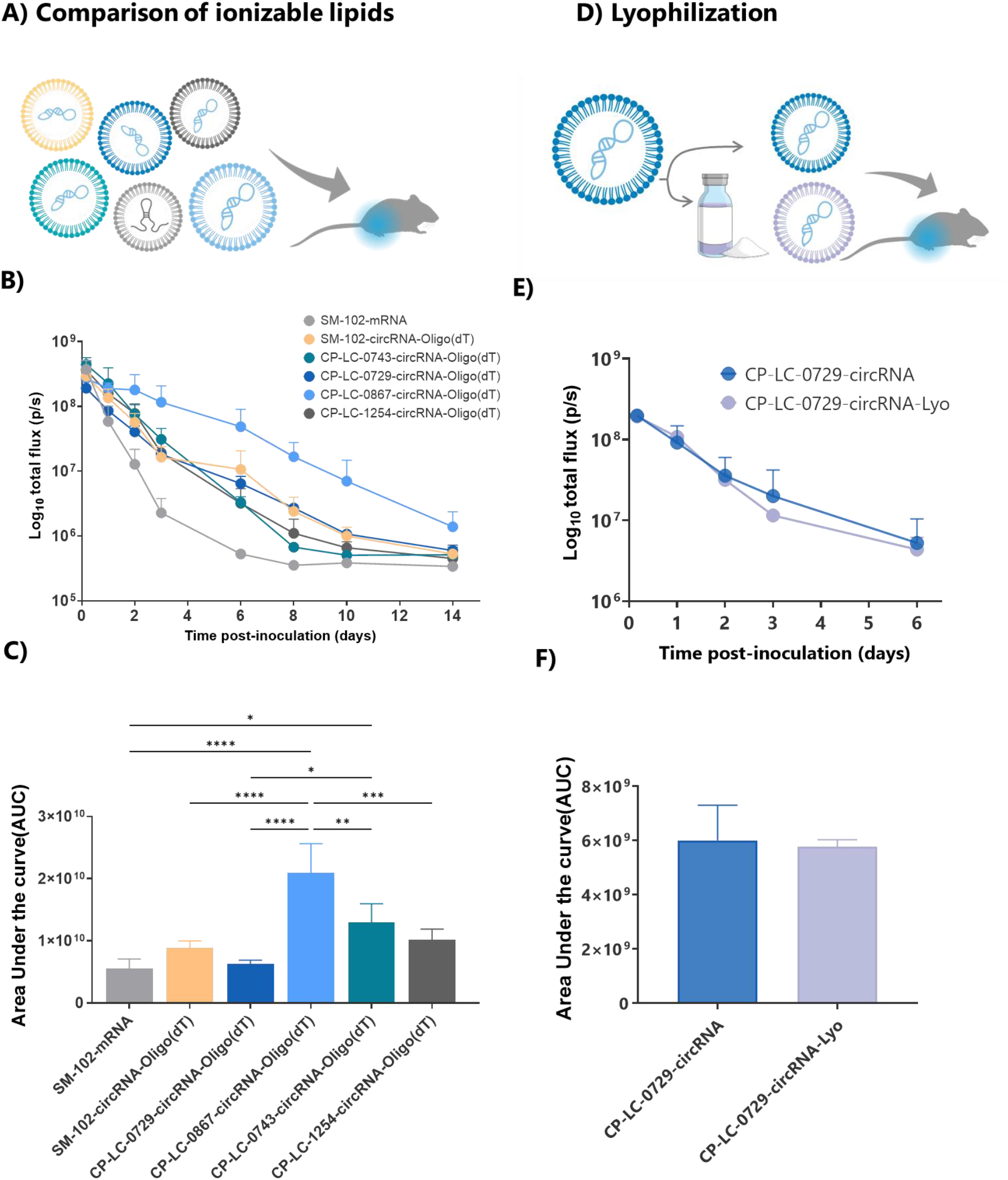
**(A)** Schematic of *in vivo* comparison of different ionizable lipids used to encapsulate circRNA-Oligo(dT) into LNPs. Control using mRNA as cargo is indicated in light grey. **(B)** Luminescence monitoring over a 14-day period following intramuscular injection in mice of LNPs encapsulating circRNA-Oligo(dT) or mRNA **(C)** Comparison of cumulative luciferase protein expression (AUC) over the entire monitoring period in panel B **(D)** Experimental workflow illustrating the lyophilization of LNPs encapsulating circRNA-Oligo(dT) using CP-LC-0729 as the ionizable lipid **(E**) Luminescence monitoring over a 6-day period following intramuscular injection in mice, comparing lyophilized and non-lyophilized LNP formulations. **(F)** Cumulative luciferase protein expression, calculated via the area under the curve (AUC) of panel E. Results are represented as mean ± SD. Statistical significance was determined using one-way ANOVA with Tukey’s post-hoc test (*: *P*-value <0.05**; *P*-value <0.01; ***: *P*-value <0.001; *absence of asterisks*: *P*-value >0.05).

Notably, while luminescence was initially planned to be monitored over a 6-day period post-injection, LNPs formulated with CP-LC-0867 exhibited sustained high levels of luciferase expression, prompting us to extend the monitoring period to 14 days. LNPs containing CP-LC-0867 consistently produced higher protein levels compared to those formulated with SM-102, while LNPs with CP-LC-0729 exhibited a similar expression profile to SM-102 (Figure 4B). CP-LC-0743 and CP-LC-1254 also showed similar expression profiles to SM-102, but with a more pronounced decrease in protein levels from 72 hours onward. Interestingly, these in vivo results did not align with the in vitro data from HeLa, HEK293T, or A549 cells (Supplementary Figure 4B), where CP-LC-0867 performed similarly to the other lipids. This discrepancy may be due to the shorter time points used in cell-based luminescence assays compared to the extended monitoring in mice. However, it is more likely attributable to the well-documented differences between in vitro and in vivo transfection efficiencies of LNPs, as reported extensively in the literature (32).

When analyzing cumulative luciferase protein expression, as measured by the area under the curve (AUC), CP-LC-0867 generated significantly higher protein levels compared to all other conditions (Figure 4C). Although the AUC analysis showed similar total luciferase production between mRNA and certain circRNA formulations, this was largely due to mRNA producing higher protein levels at early time points, which skews the overall comparison. However, at an intermediate time point of 6 days, the sustained protein production profile became more evident, with CP-LC-0867 exhibiting a 92-fold increase in protein expression and the remaining circRNA formulations showing at least a 6-fold increase compared to mRNA-LNP formulations. These results highlight the long-term stability and effectiveness of circRNA-LNPs, particularly those formulated with CP-LC-0867.

We then assessed the lyophilization capacity of LNP-circRNA-Oligo(dT) to extend its storage life and eliminate the need for ultra-low temperature storage and transport, which represents a significant hurdle to the broader adoption of circRNA therapies. For this study, we selected CP-LC-0729 as the ionizable lipid, as it is one of the most thoroughly characterized lipids among the new ionizable lipids we have tested. Using a previously established lyophilization method (24), we lyophilized LNPs encapsulating circRNA-Oligo(dT) with CP-LC-0729 and evaluated their physical characteristics (Supplementary Figure 4) and functionality in mice, comparing them with non-lyophilized LNPs (Figure 4D). After intramuscular administration and monitoring luminescence over a 6-day period, we observed no significant differences in the protein production profile (Figure 4E) or cumulative protein expression (Figure 4F) between lyophilized and non-lyophilized LNPs.

These results indicate that lyophilization does not impair the functionality of LNP-circRNA-Oligo(dT) using CP-LC-0729 as ionizable lipid, making this new platform a viable option for extending storage life without the need for ultra-low temperature conditions.

## DISCUSSION

Circular RNA (circRNA) has gained significant attention in therapeutic applications due to its exceptional stability and resistance to exonucleases. However, achieving efficient and scalable purification remains a considerable challenge. Traditional methods, such as PAGE electrophoresis and RNase R digestion, while effective, present notable limitations regarding throughput, scalability, and purity (33). PAGE, commonly employed for RNA separation based on size, conformation and total molecular charge, provides high resolution but is labor-intensive and prone to contamination, which significantly reduces its practicality for large-scale operations (34). Likewise, RNase R digestion, which selectively degrades linear RNA while preserving circRNA, often leaves structured linear contaminants and proves cost-prohibitive for large-scale purification. Consequently, while these methods are suitable for initial characterization, they fall short in terms of scalability and widespread industrial application (35).

High-performance liquid chromatography (HPLC) has also been explored for its potential to remove impurities such as double-stranded RNA (dsRNA) (9). However, the HPLC technique reported for circRNA suffer from low recovery yields, as only a small fraction of the total RNA is collected, thus making the method challenging to scale for industrial use (9). In contrast, our optimized affinity chromatography method, which employs oligo(dT) columns, effectively addresses these issues by significantly enhancing the yield of circRNA, while being compatible with scalable processes.

In relation to Mg^2+^ concentration during the IVT process, previous reports largely rely on commercial kits where Mg^2+^ concentrations are undisclosed, making it impossible to modify or optimize (9, 11, 13). In this study, we demonstrated that by adjusting the Mg^2+^ concentration, we can significantly increase the proportion of circRNA produced, providing an important optimization strategy for circRNA synthesis.

In terms of *in vivo* delivery of circRNA, only a limited number of studies have reported successful strategies. Some approaches have employed cationic polymers such as *in vivo*-jetPEI (11) or O6-stat-N6 CARTs (3, 6), while others have focused on LNPs using ionizable lipids like cKK-E12 (11), MC3 (2, 36), SM-102, and ALC-0315 (28). Nonetheless, several studies that utilize LNPs do not specify the ionizable lipid used (19, 22). Here we report the best-performing ionizable lipid described to date, which demonstrated a remarkable 92-fold increase in *in vivo* protein expression at 6 days post-injection compared to SM-102. Moreover, while previous studies typically employed 5 to 20 μg of RNA for *in vivo* delivery (2, 3, 21, 27, 36), we achieved robust protein expression with just 1 μg of circRNA. This demonstrates the exceptional efficiency of our novel lipid formulations and their potential to significantly reduce the required therapeutic circRNA dosage.

A critical factor in evaluating circRNA for clinical applications is the duration of gene expression, as circRNA is designed to provide more sustained and stable expression compared to mRNA. To our knowledge, the longest previously reported *in vivo* signal duration is 42 hours using LNPs (13) and 168 hours using O6-stat-N6 CARTs (6). In this study, we report an extended signal duration of 336 hours, significantly surpassing these previous results. This prolonged expression suggests that the combination of Oligo(dT) purification and novel ionizable lipids provides superior performance over existing methods, making this approach particularly promising for long-term therapeutic applications.

Lyophilization has been extensively studied to enhance the stability of mRNA vaccines (24), but there has been limited research on its application to circRNA-LNPs, with only one study available on the topic (21). However, this study does not specify the ionizable lipid used and focuses on mannose-modified PEG for circRNA-LNP delivery to lymphocytes, without assessing the impact of lyophilization on LNPs with unmodified PEG. In our study, we evaluated the effects of lyophilization on conventional PEGylated LNPs using a new ionizable lipid (22) encapsulating circRNA, offering a more comprehensive analysis of lyophilization’s potential in maintaining the physical integrity and functionality of circRNA-LNP formulations.

Despite the promising results, our study has some limitations that warrant further investigation. First, while we have demonstrated enhanced circRNA yield and prolonged *in vivo* expression using Oligo(dT) purification and novel ionizable lipids, the evaluation was based on intramuscular injection This route of administration is widely used for vaccines and other therapeutic applications, providing valuable insights into the platform’s effectiveness. However, it may not fully capture how the system would perform with alternative delivery routes, such as intravenous or subcutaneous injections. Exploring these routes in future studies will offer a broader understanding of the platform’s therapeutic potential and expand its applicability across various clinical scenarios. Additionally, we successfully lyophilized circRNA-LNPs, preserving their functionality, and developed an optimized and streamlined process for generating high-quality circRNA. Together, these advancements mark a qualitative breakthrough, setting the stage for more efficient and scalable circRNA therapeutic development. Looking ahead, we aim to explore the long-term stability of lyophilized circRNA-LNPs, building on our previous success with mRNA-LNPs stored for over a year at 4ºC (24), which included the use of CP-LC-0867 and other top-performing lipids. In addition, we will delve deeper into our extensive library of over 1,500 ionizable lipids to identify even more effective candidates for circRNA delivery. By deciphering the underlying mechanisms driving LNP efficacy, we will push the boundaries of circRNA delivery, bringing this modality closer to widespread clinical application.

In summary, our study provides an important step forward in the scalable production and *in vivo* delivery of circular RNA (circRNA), utilizing optimized purification strategies and novel ionizable lipids. We demonstrated that the use of Oligo(dT) affinity chromatography enhances the yield and purity of circRNA, allowing for efficient downstream applications without the need for additional enzymatic treatments. Furthermore, we successfully encapsulated circRNA into LNPs formulated with STAAR of ionizable lipids, including CP-LC-0867 and CP-LC-0729, which resulted in superior protein expression profiles compared to conventional lipids such as SM-102. Additionally, our exploration of lyophilization revealed that lyophilized circRNA-LNP formulations retain their functionality, featuring their potential for long-term storage and global distribution. These results demonstrate the substantial potential of this platform to advance the clinical translation of circRNA therapies by enhancing production efficiency, stability, and delivery, which are critical factors for achieving reliable and scalable therapeutic outcomes.

## DATA AVAILABILITY

Data available on request.

## SUPPLEMENTARY DATA

## AUTHOR CONTRIBUTIONS

Verónica Lampaya: Conceptualization, Data curation, Formal analysis, Investigation, Methodology, Validation, Visualization, Writing – review & editing. Ana Larraga: Conceptualization, Data curation, Formal analysis, Investigation, Methodology, Validation, Visualization, Writing – review & editing. Víctor Navarro: Investigation and Methodology. Alexandre López: Data curation, Formal analysis, Investigation, Methodology, Validation, Visualization, Writing -review & editing. Diego de Miguel: Conceptualization, Supervision and Writing – review & editing. Álvaro Peña: Investigation, Methodology and Writing – review & editing. Esther Broset: Conceptualization, Formal analysis, Supervision, Visualization, Writing – original draft. Juan Martínez-Oliván: Conceptualization, Funding acquisition, Supervision, Writing – review & editing. Diego Casabona: Conceptualization, Project administration, Supervision and Writing – review & editing.

## ACKNOWLEDGEMENTS

We would like to extend our sincere gratitude to all the members of Certest Biotec, especially the Certest Pharma group, whose valuable insights and expertise significantly contributed to the success of this research. Authors would like to acknowledge the use of “Servicios Científico Técnicos” at CIBA (IACS-University of Zaragoza).

