## Supplementary for "A complete approach for circRNA therapeutics from purification to lyophilized delivery using novel ionizable lipids"

#### SUPPLEMENTARY FIGURES

Supplementary Figure 1

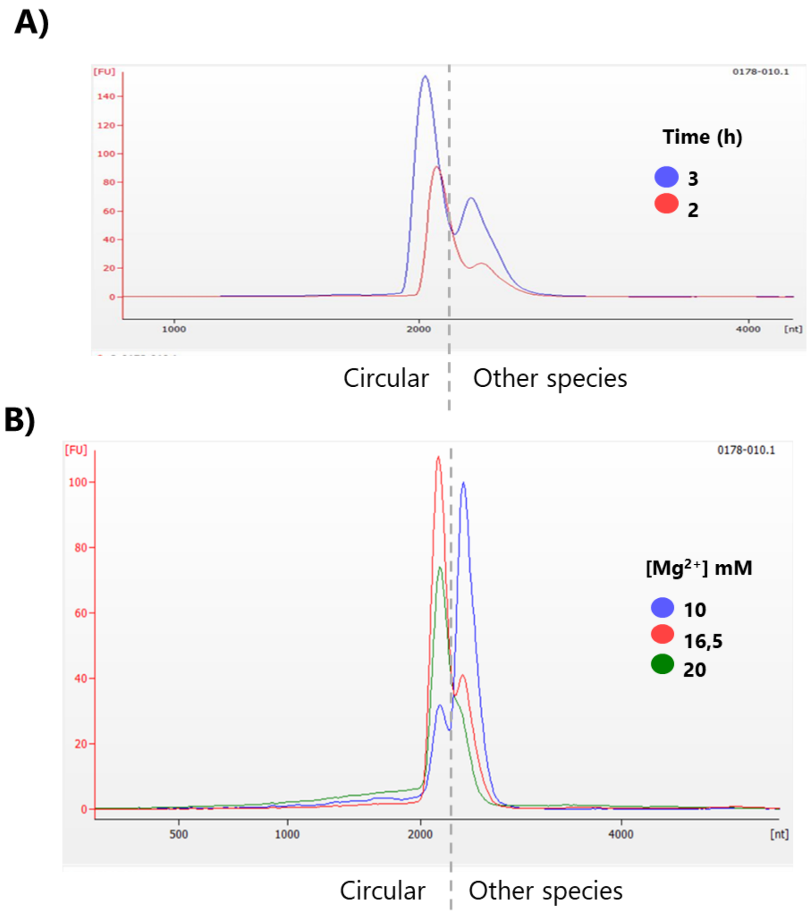


**Supplementary Figure 1.** Capillary electrophoresis histograms **(A)** at various incubation times during the in vitro transcription (IVT) reaction optimization and **(B)** with various $Mg^{2+}$ concentrations. The left peak corresponds to the circular RNA isoforms, while the right peak corresponds to the remaining RNA species.

Supplementary Figure 2

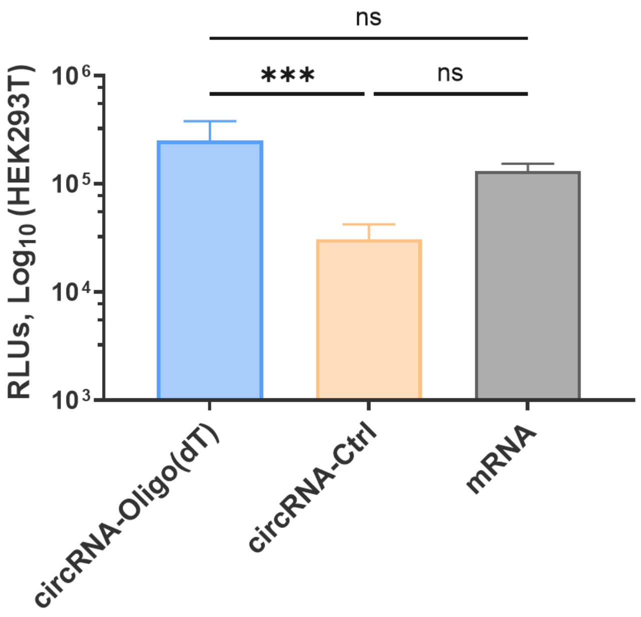


**Supplementary Figure 2.** Luciferase production measured in Relative Luminescence Units (RLU) in HEL293T cell line transfected with 100 ng/well of circRNA-Oligo(dT), circRNA-Ctrl or mRNA. Luminescence was measured 24 hours post-transfection. Results are represented as mean ± SD. Statistical significance was determined using one-way ANOVA with Tukey's post-hoc test (***: *P*-value <0.001; *ns*: *P*-value >0.05).

#### Supplementary Figure 3

CP-LC-0743

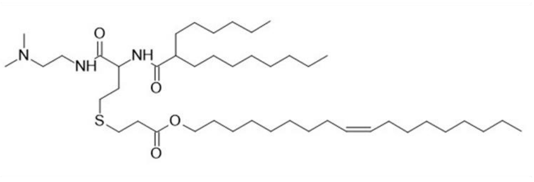

CP-LC-0729

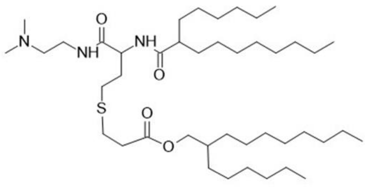

CP-LC-1254

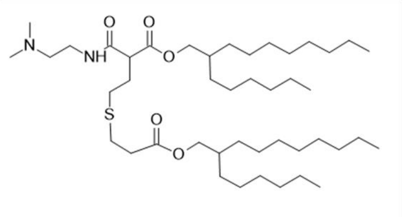

CP-LC-0867

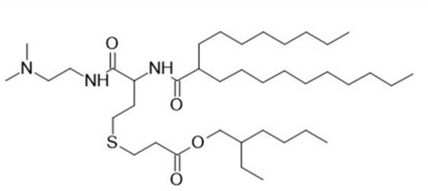

**Supplementary Figure 3. Structure of STAAR ionizable lipids used in the study.**

### Supplementary Figure 4

**A)**

| RNA type | Ionizable lipid | Dimeter [nm] | Polydispersity index | Zeta Potential [mV] | Encapsulation % |
|---|---|---|---|---|---|
| mRNA | SM-102 | 98,57 | 0,2325 | -5,164 | 90,69 |
| circRNA-oligo(dT) | SM-102 | 109,9 | 0,2318 | -7,938 | 93,38 |
| | CP-LC-0729 | 89,93 | 0,2912 | -11,03 | 90,47 |
| | CP-LC-0867 | 113,3 | 0,2504 | -8,672 | 84,68 |
| | CP-LC-0743 | 136,9 | 0,06148 | -4,859 | 83,46 |
| | CP-LC-1254 | 107,8 | 0,2943 | -9,782 | 93,98 |

**B)**

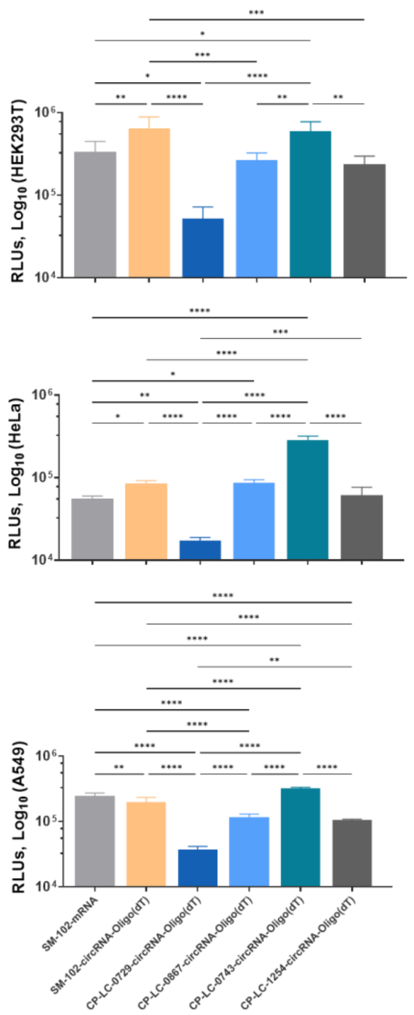


**Supplementary Figure 4. (A)** Table summarizing the physical properties of LNP formulations containing circRNA or mRNA and the indicated ionizable lipid. The measured parameters include particle diameter (in nanometers), polydispersity index (PDI) for uniformity, zeta potential (in mV) for surface charge, and encapsulation efficiency (percentage of RNA encapsulated). **(B)** Luciferase production measured in Relative Luminescence Units (RLU) in HEK293T, HeLa and A549 cell lines transfected with 100 ng/well of LNPs encapsulating circRNA-Oligo(dT) or mRNA with the indicated ionizable lipids. Luminescence was measured 24 hours post-transfection. Results are represented as mean ± SD. Statistical significance was determined using one-way ANOVA with Tukey's post-hoc test (*: P-value <0.05**; P-value <0.01; ***: P-value <0.001; ****: P-value <0.0001 ns: P-value >0.05).

### Supplementary Table 5

| RNA type | Formulation | Ionizable lipid | Dimeter [nm] | Polydispersity index | Zeta Potential [mV] | Encapsulation % |
|---|---|---|---|---|---|---|
| circRNA-oligo(dT) | Liquid | CP-LC-0729 | 92,84 | 0,2342 | -14,49 | 96,10 |
| circRNA-oligo(dT) | Lyophilized | CP-LC-0729 | 101,9 | 0,1552 | -11,8 | 86,20 |

**Supplementary Figure 5.** Table summarizing the physical properties of LNP formulations containing circRNA-Oligo(dT) and CP-LC-0729 ionizable lipid either lyophilized or not. The measured parameters include particle diameter (in nanometers), polydispersity index (PDI) for uniformity, zeta potential (in mV) for surface charge, and encapsulation efficiency (percentage of RNA encapsulated).
